# 7q11.23 Syndromes Reveal BAZ1B as a Master Regulator of the Modern Human Face and Validate the Self-Domestication Hypothesis

**DOI:** 10.1101/570036

**Authors:** Matteo Zanella, Alessandro Vitriolo, Alejandro Andirko, Pedro Tiago Martins, Stefanie Sturm, Thomas O’Rourke, Magdalena Laugsch, Natascia Malerba, Adrianos Skaros, Sebastiano Trattaro, Pierre-Luc Germain, Giuseppe Merla, Alvaro Rada-Iglesias, Cedric Boeckx, Giuseppe Testa

**Affiliations:** Department of Oncology and Hemato-Oncology, University of Milan, Italy; Department of Experimental Oncology, Laboratory of Stem Cell Epigenetics, European Institute of Oncology, Milan, Italy; University of Barcelona, Spain; University of Barcelona, Institute of Complex Systems, Barcelona, Spain; Center for Molecular Medicine Cologne (CMMC), University of Cologne, Germany; Institute of Human Genetics, CMMC, University Hospital, Cologne, Germany; Division of Medical Genetics, Fondazione IRCCS Casa Sollievo della Sofferenza, San Giovanni Rotondo, Foggia, Italy; Current address: Brain Research Institute, University of Zürich, Switzerland; Institute of Biomedicine and Biotechnology of Cantabria (IBBTEC), University of Cantabria, Spain; Institute for Advanced Studies and Research (ICREA), Barcelona, Spain

**Author notes:** These authors contributed equally to this work (Matteo Zanella, Alessandro Vitriolo).

## Abstract

Symmetrical 7q11.23 dosage alterations cause craniofacial and cognitive/behavioral phenotypes that provide a privileged entry point into the evolution of the modern human face and (pro-) sociality. We undertook a functional dissection of chromatin remodeler BAZ1B in neural crest stem cells (NCSCs) from a uniquely informative cohort of typical and atypical patients harboring 7q11.23 Copy Number Variants (CNVs). Our results reveal a key contribution of BAZ1B to NCSC *in vitro* induction and migration, coupled with a crucial involvement in neural crest (NC)-specific transcriptional circuits and distal regulation. By intersecting our experimental data with new paleogenetic analyses comparing modern and archaic humans, we uncover a modern-specific enrichment for regulatory changes both in BAZ1B and its experimentally defined downstream targets, thereby providing the first empirical validation of the self-domestication hypothesis and positioning BAZ1B as a master regulator of the modern human face.

**One Sentence Summary:** BAZ1B dosage shapes the modern human face.

---

Anatomically modern humans (AMHs) exhibit a suite of craniofacial and prosocial characteristics that are reminiscent of traits distinguishing domesticated species from their wild counterparts (*1*, *2*). Accordingly, the self-domestication hypothesis predicts that key aspects of modern humans’ anatomy and cognition can be illuminated by studies of the so called "domestication syndrome", the core set of domestication-related traits that was recently proposed to result from mild neural crest (NC) deficits (*3*). Both the neurocristopathic basis of domestication and its extension to the evolution of AMHs remain however to be tested experimentally. Williams-Beuren syndrome (WBS, OMIM 194050) and Williams-Beuren region duplication syndrome (7dupASD, OMIM 609757), caused respectively by the hemideletion or hemiduplication of 28 genes at the 7q11.23 region (WBS critical region, hereafter WBSCR), represent a paradigmatic pair of neurodevelopmental conditions whose NC-related craniofacial dysmorphisms (Fig. S1A) and cognitive/behavioral traits (*4*, *5*) are uniquely and symmetrically aligned to the domestication-related traits of AMHs. Structural variants in WBS genes have been shown to underlie stereotypical hypersociability in domestic dogs (*6*) and within the WBSCR the following lines of evidence point to the chromatin regulator *BAZ1B* as a key gene underlying domestication-relevant craniofacial features: i) its crucial role in NC maintenance and migration in *Xenopus Laevis* and the craniofacial defects observed in knock-out mice (*7*, *8*); ii) the observation that its expression is impacted by domestication-related events in canines (*9*); iii) the first formulation of the neurocristopathic hypothesis of domestication, which included *BAZ1B* among the genes influencing NC development (*3*); iv) the most comprehensive studies focusing on regions of the modern human genome associated with selective sweep signals compared to Neanderthals/Denisovans (hereafter ‘archaics’) (*10*, *11*), one of which specifically included *BAZ1B* within the detected portions of the WBSCR; and v) the thus far most detailed study systematically exploring high frequency (> 90%) changes in modern humans for which archaic humans carry the ancestral state, which found *BAZ1B* enriched for mutations in modern humans (most of which fall in the regulatory regions of the gene) (*12*). Through the thus far largest cohort of 7q11.23 patient-derived induced pluripotent stem cell (iPSC) lines, our previous work had revealed major disease-relevant dysregulation that was already apparent at the pluripotent state and was further exacerbated upon differentiation (*13*). Here we build on this insight to specifically dissect the impact of BAZ1B dosage alterations on the NC of WBS and 7dupASD patients and to empirically test the self-domestication hypothesis by interrogating the correlative evidence from paleogenomic analyses with the experimentally determined BAZ1B-dependent circuits underlying craniofacial morphogenesis. We demonstrate a major contribution of BAZ1B to the modern human face, validating the prediction that posits the evolution of the human face as an instance of mild neurocristopathy.

## Results

### Establishment and validation of an extensive cohort of patient-specific BAZ1B-interfered NCSC lines

In order to dissect the role of BAZ1B in the paradigmatic craniofacial dysmorphisms that characterize WBS and 7dupASD, we started from our previous characterization of WBS and 7dupASD patient-specific iPSC lines and differentiated derivatives (*13*) and selected a cohort of 11 NCSC lines (4 from WBS patients, 3 from 7dupASD patients and 4 from control individuals). These represent the largest cohort of patient-specific NCSCs described so far and we used transcriptomic profiling to validate their cranial identity as a privileged entry point into disease-pathogenesis, given the centrality of the cranial NC in the development of the face. We then knocked-down (KD) BAZ1B via RNA interference in all lines across the three genetic conditions, including also NCSCs derived from a particularly informative atypical WBS patient (hereafter atWBS) bearing a partial deletion of the region that spares *BAZ1B* and six additional genes (*14*) (Fig. 1A). In order to establish a high-resolution gradient of BAZ1B dosages, we selected two distinct shRNA against BAZ1B (*i.e.*, sh1 and sh2) along with a scrambled shRNA sequence (hereafter scr) as negative control, for a total of 32 NCSC lines. KD efficiency was evaluated both at the RNA level by quantitative PCR (qPCR) (Fig. 1B; S1C), confirming the attainment of the desired gradient with an overall reduction of about 40% for sh1 and 70% for sh2, as well as reduction at the protein level, as detected by Western blot (Fig. S1E).

**Figure 1.**
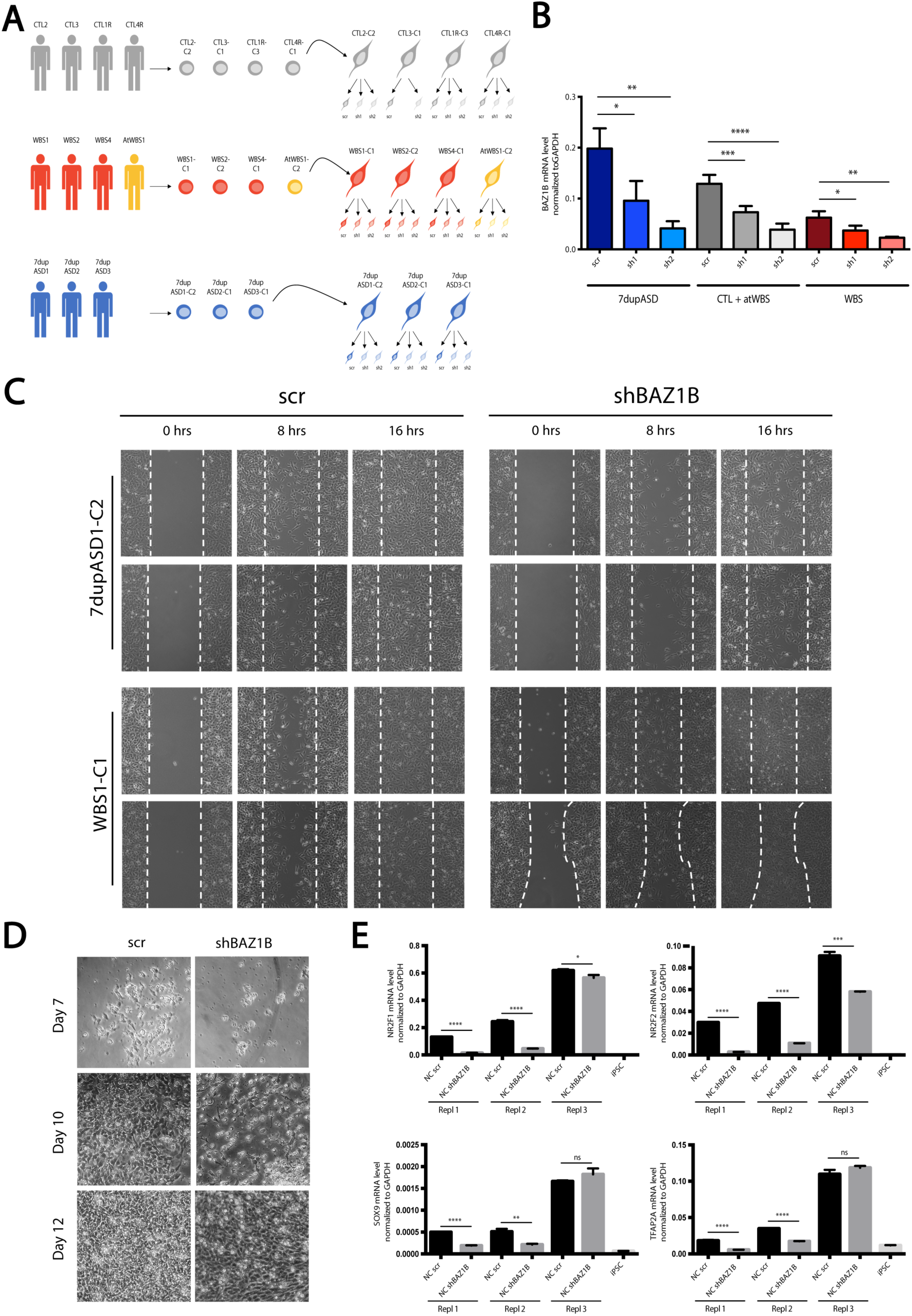
BAZ1B KD impairs migration and induction of patient-specific iPSC-derived NCSCs. (**A**) Schematic representation of the KD strategy on our iPSC-derived NCSC cohort. (**B**) BAZ1B mRNA levels in all the interfered lines (scr, sh1 and sh2) as measured by qPCR. Data represent aggregates of samples with the same number of BAZ1B copies (7dup, CTL + atWBS and WBS). GAPDH is used as normalizer. (**C**) 8 hours and 16 hours-time points from the wound-healing assay analysis performed on a 7dupASD and a WBS NCSC line upon BAZ1B KD. Cells from the same line infected with the scr sh were used as references for the migration (n=2). (**D**) Day 7, 10 and 12 of NC differentiation from EBs of a scr-interfered iPSC line and its respective BAZ1B KD (n=3). (**E**) mRNA levels of NC markers at day 12 of differentiation (experiment reported in panel D). An iPSC line is included as a negative control. Student’s t-test (ns: non-significant; *: p-value < 0.05; **: p-value < 0.01; ***: p-value < 0.001 and ****: p-value < 0.0001).

### BAZ1B interference impairs NCSC migration and induction

NCSCs need to migrate in order to reach specific target regions in the developing embryo and give rise to distinct cell types and tissues, including craniofacial structures. BAZ1B KD was shown to affect the migration but not the induction of the NC in *Xenopus laevis,* and to promote cancer cell invasion in different lung cancer cell lines (*7*, *15*). We thus hypothesized that the *BAZ1B* dosage imbalances entailed in the 7q11.23 syndromes could result in a defective regulation of NCSC migration and might hence underlie the NC-related alterations typical of WBS and 7dupASD patients. To test this, we compared the migration properties of patient-specific BAZ1B KD NCSC lines (sh2) to their respective control NCSC line (scr) by the well-established wound-healing assay. The 7dupASD NCSC KD lines took longer to fill the wound when compared to the respective control lines (scr), as indicated by images taken at 8 and 16 hours after a gap was created on the plate surface (Fig. 1C, S1F). We instead observed an opposite behavior for the WBS BAZ1B KD lines, which were faster than the respective scr lines in closing the gap (Fig. 1C, S1F). Interestingly, in contrast to the previous observations from *Xenopus laevis* (*7*), we also observed a minor delay in NC induction as a consequence of BAZ1B KD (Fig. S1D), by means of a differentiation protocol based on NC delamination from adherent embryoid bodies (EBs), which recapitulates the initial steps of NC generation (*16*). In particular, starting from 2/3 days after attachment of EBs, we observed a lower number of outgrowing cells in the KD line (Fig. 1D, day 7 and 10), coupled with an evidently higher cell mortality. Cells were eventually able to acquire the typical NC morphology, even though lower differentiation efficiency was evident, as shown by images taken at day 12. In addition, the delay in NC formation was associated with a downregulation of well-established critical regulators of NC migration and maintenance, including *NR2F1*, *NR2F2*, *TFAP2A* and *SOX9* (Fig. 1E). These findings show that BAZ1B regulates the developing NC starting from its earliest migratory stages and that the symmetrically opposite 7q11.23 dosages alterations prime NCSCs to symmetrically opposite deficits upon BAZ1B interference.

### BAZ1B interference disrupts key neural crest-specific transcriptional circuits

A core insight from our previous work had been the discovery that already at the pluripotent state 7q11.23 CNVs cause transcriptional dysregulation in disease-relevant pathways, which are then selectively amplified along development in a tissue-specific manner (*13*). Given the critical role of BAZ1B as transcriptional regulator in different cell and animal models (*17*–*19*), as well as its functional involvement in human NC from the above described findings, we thus hypothesized that its dosage could significantly contribute to the transcriptional dysregulation in 7q11.23 CNV-NCSCs. To test this hypothesis and gain mechanistic insights into the specific BAZ1B dosage-dependent downstream circuits, we subjected 32 interfered NCSC lines to high coverage RNA sequencing (RNA-seq) analysis. As shown in Fig. S2A, a manually curated signature from an extensive literature review (*20*–*25*) validated the cranial identity of our NCSC lines, while clustering by Pearson correlation excluded the presence of any genotype-or hairpin-specific expression change. Confirming our previous observations in the two largest cohorts of iPSC lines (*26*), a Principal Component Analysis (PCA) corroborated the significant impact of individual genetic backgrounds on transcriptional variability, with most ‘KD lines’ clustering with their respective control ‘scr line’. This was consistent with the narrow range of experimentally interfered BAZ1B dosages and pointed to a selective BAZ1B dosage-dependent transcriptional vulnerability (Fig. S2B).

In order to dissect it, we thus resorted to a combination of classical pairwise comparative analysis, contrasting shBAZ1B-interfered NCSC lines (sh1+sh2) with their respective controls (scr), with a complementary regression analysis using *BAZ1B* expression levels as independent variables, subtracting the contribution of individual genetic backgrounds. This design increases robustness and sensitivity in the identification of genes that, across multiple genetic backgrounds and target gene dosages, might have a different baseline (scr) across individuals while still being robustly dysregulated upon BAZ1B interference.

The two analyses identified a total of 448 genes with FDR < 0.1 (1192 with p <0.01 and FDR < 0.25) whose transcriptional levels followed BAZ1B dosage, either in a direct (202; 539 with p < 0.01 and FDR < 0.25) or inverse (246; 653 with p < 0.01 and FDR < 0.25) fashion. In addition, genes identified only in the regression analysis included around 90% of the differentially expressed genes (DEGs) (27/29, FDR < 0.1) found in the comparative analysis (Fig. 2A). Consistent with the differential efficiency of the two short hairpins, we found a globally stronger transcriptional impact for the group of samples targeted by sh2 (Fig. S2C) and a milder but nevertheless clearly distinguishable effect of sh1, resulting in particularly informative gradient of dosages over the scr control lines.

**Figure 2.**
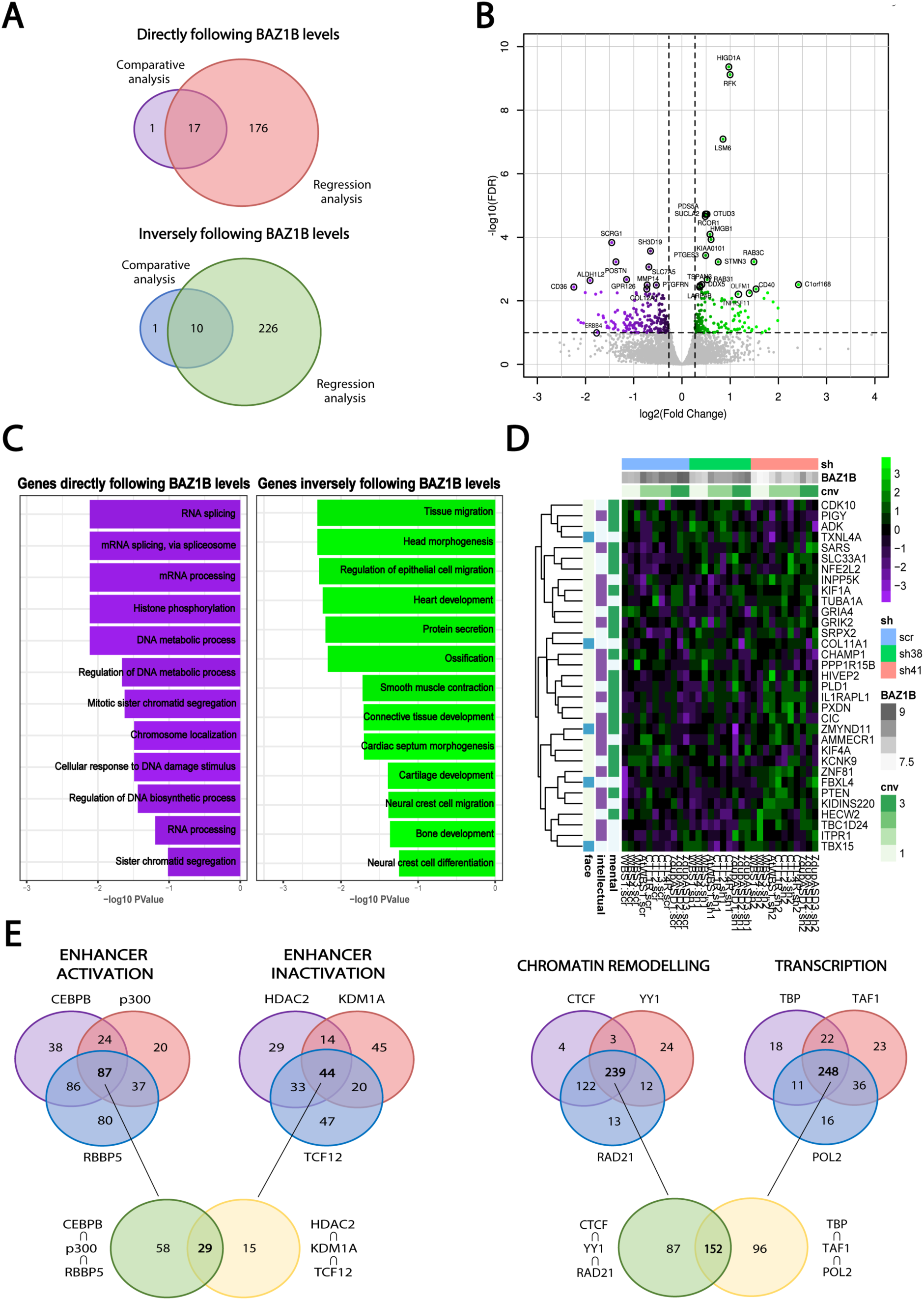
BAZ1B KD is responsible for transcriptional alterations in NC-related pathways. (**A**) Overlap between genes directly or inversely following BAZ1B levels identified in the pairwise comparative analysis (scr *vs* shBAZ1B) and in the regression analysis on BAZ1B-level sensitive genes on iPSC-derived NCSCs (FDR < 0.1). (**B**) Volcano plot reporting DEGs identified in the RNA-seq analysis on iPSC-derived NCSCs (FC > 1.25; FDR < 0.1). (**C**) Top most-specific enrichments for GO biological processes among the DEGs in the RNA-seq analysis on iPSC-derived NCSCs. (**D**) Heatmap representing DEGs that are dysregulated in genetic disorders involving mental retardation, intellectual disability and/or facial dysmorphisms according to OMIM database classification. (**E**) Putative regulators of genes that follow BAZ1B levels identified by a Master regulator analysis. Regulators were divided in four different groups based on their main functions.

Particularly noteworthy among the genes that we found inversely correlated with *BAZ1B* levels by highest fold changes (FC > 2) were the following crucial regulators of cranial NC: i) *POSTN*, which plays a key role in cranial NC-mediated soft palate development (*27*) and is responsible for heart valve defects and periodontal disease-like phenotypes in KO mice (*28*); and ii) *ERBB4* which is important for skeletal muscle development and NC migration (*29*) and causes heart defects and aberrant cranial NC migration in deficient mice (*30*) (Fig. 2B). Genes that instead directly follow BAZ1B levels include: i) *OLFM1,* whose overexpression promotes an increased and continued production of NC cells (*31*); ii) *TNFRSF11B,* involved in bone resorption and osteoclast activation (*32*) (Fig. 2B); iii) *CUL3*, which has been involved in craniofacial morphogenesis and ubiquitinates TCOF1 and NOLC1 as a master regulator of NC specification (*33*); and iv) the CUL3 adaptor *KLHL12* (*34*). As both *KLHL12* and *TCOF1*, acting up-and down-stream of *CUL3* respectively, are involved in craniofacial morphogenesis (*35*), these data highlight a convergent BAZ1B dosage-dependent dysregulation of the foundational *CUL3*-centered regulatory axis orchestrating NC-mediated craniofacial morphogenesis.

Additional particularly relevant genes inversely following BAZ1B dosage are: i) *TBX15*, one of the recently mapped loci associated to the variability of human facial shape (*36*); ii) *AMMECR1*, whose mutations cause midface hypoplasia and hearing impairment (*37*); iii) *ZMYND11*, a H3K36me3 reader that, when mutated, gives rise to dysmorphic facial features and mild intellectual disability (*38*, *39*); iv) *SNAI2*, a critical regulator of NC-based organogenesis (*40*); and v) *BBC3*, which is repressed by *SNAI2* in response to DNA damage (*41*).

GO analysis performed on genes directly following BAZ1B levels suggested specific enrichments in biological process such as histone phosphorylation, chromosome localization, RNA processing and splicing. Genes inversely following BAZ1B levels were instead enriched in categories particularly relevant for NC and NC-derivative functions, such as cell migration and cardiovascular and skeletal development (Fig. 2C). Interestingly, by interrogating the OMIM database, we found that several DEGs were associated with genetic disorders whose phenotypes include “mental retardation”, “intellectual disability” and/or “facial dysmorphisms” (Fig. 2D), underscoring the pertinence of BAZ1B-dependent dysregulation across both the neurocristopathic and cognitive axes.

Finally, a Master Regulator Analysis identified candidate regulators of BAZ1B DEGs, including factors involved in enhancer marking (CEBPB, p300, RBBP5, HDAC2, KDM1A and TCF12), promoter activation (TBP, TAF1 and POL2) and chromatin remodeling (CTCF, RAD21 YY1) (Fig. 2E; S2C), several of which are themselves causative genes of intellectual disability (ID) syndromes with neurocristopathic involvement, as in the case of our recently identified Gabriele-de Vries syndrome caused by YY1 haploinsufficiency (*42*, *43*). Chromatin remodeling was indeed the most prominently enriched group within the overall domain of transcriptional regulation. Two master regulators are particularly noteworthy as they are themselves regulated by BAZ1B dosage. The first is EGR1 (FDR < 0.1), itself among the genes inversely correlated with BAZ1B levels, which is implicated in cranial development (in animal models) (*44*, *45*) and whose promoter has been recently shown to feature a bivalent state in human embryonic cranial NC (*20*, *33*). The second is MXI1, identified as master regulator of genes directly following BAZ1B levels (FDR < 0.001), which is itself found among the genes inversely correlated with BAZ1B and is itself a regulator of BAZ1B, pointing to a crosstalk between the two (Fig. S2C). Notably, two differentially expressed targets of MXI1, *TGFB2* and *NFIB,* are also known to be involved in intellectual disability and craniofacial defects (*46*–*48*).

### BAZ1B regulates the neural crest epigenome in a dosage-dependent manner

In order to mechanistically anchor transcriptional dysregulation to the dosage-dependent occupancy of BAZ1B, we set out to define its direct targets and investigate their chromatin state at both promoters and enhancers upon BAZ1B KD by chromatin immunoprecipitation coupled with sequencing (ChIP-seq). Given the absence of ChIP-grade antibodies, we designed a tagging strategy to establish, by CRISPR/Cas9 editing, a series of in-frame 3xFLAG endogenously tagged BAZ1B alleles in representative iPSCs of the four genotypes (Fig. 3A and S3A, B). These were then differentiated to NCSCs (Fig. S3C) and subjected to ChIP-seq with anti-FLAG antibody, enabling a faithful characterization of BAZ1B genome-wide occupancy across dosages (one tagged allele in WBS, two tagged alleles in atWBS and CTL, and two tagged alleles in the context of 1.5-fold dosages in 7dupASD).

**Figure 3.**
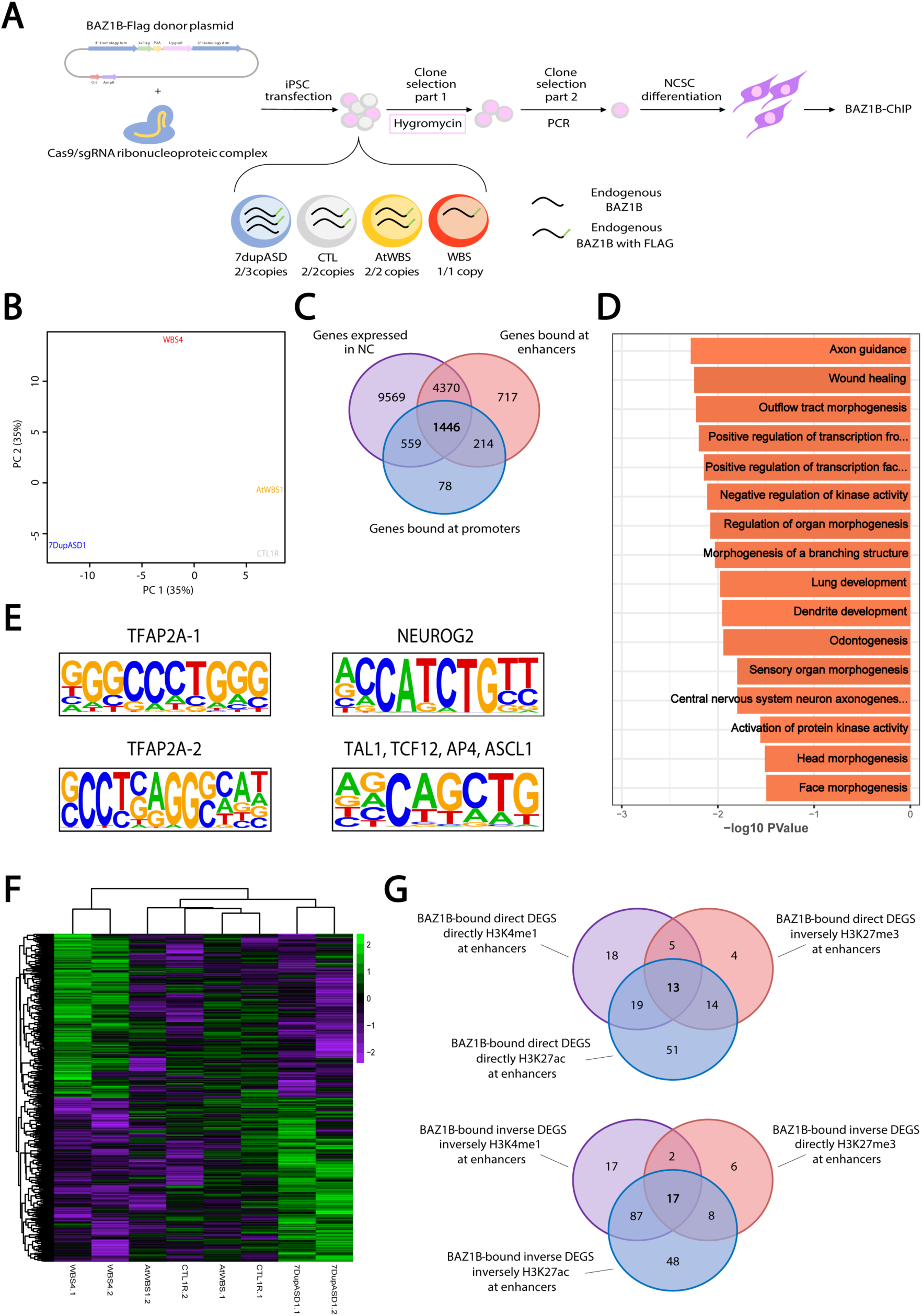
BAZ1B preferentially binds its targets at their enhancer regions and its KD causes a redistribution of enhancer histone marks. (**A**) Schematic representation of the strategy for CRISPR/Cas9-mediated tagging of endogenous BAZ1B. Briefly, iPSCs from the four genotypes were electroporated with the donor plasmid and the Cas9/single guide RNA (sgRNA) ribonucleoprotein complex, clones were selected via Hygromycin and PCR, differentiated to NCSCs and then subjected to ChIP-seq. (**B**) Principal component analysis showing the distribution of the four BAZ1B-tagged NCSC lines according to their chromatin profiles. (**C**) Overlap between genes expressed in our NCSC lines (purple) and genes bound by BAZ1B at their enhancer (red) or promoter (blue) regions. (**D**) Top most-specific enrichments for GO biological processes among the genes that are bound by BAZ1B and expressed in our NCSC cohort. (**E**) Most represented BAZ1B DNA-binding motifs identified by HOMER show high similarity to neural and NC stem cell specific transcription factors motifs. (**F**) BAZ1B differentially bound regions according to its copy number (FDR < 0.1, n=2). (**G**) Overlap between genes that are differentially expressed, have their enhancers differentially marked concordantly (H3K27ac, H3K4me1 and H3K27me3) and are bound by BAZ1B at enhancers.

PCA shows a clear separation of the samples by BAZ1B copy-number, with CTL and atWBS samples clustering more closely and WBS and 7dupASD samples clustering at opposing positions (Fig. 3B). To call NC-specific enhancer regions and promoter-enhancer associations we exploited the unprecedented resolution afforded by the patients’ cohort with its underlying variability in chromatin annotation and proceeded to: i) select chromosomal regions featuring H3K4me1 and H3K27ac in at least two individuals; ii) exclude regions marked with H3K4me3 in at least two individuals; iii) eliminate regions bearing a Transcription Start Site (TSS); and iv) associate each putative enhancer to the closest TSS, identifying a total of 308470 putative enhancer regions. Notably, BAZ1B binds 75% of its targets at their enhancer regions (6747 genes), with the remaining 2297 targets bound at promoters (Fig. 3C). In addition, 40% of genes expressed in NC are bound by BAZ1B, either exclusively at enhancers (27.4%), or exclusively at promoters (3.5%) or at both regions (9%). This highlights its pervasiveness within the NC epigenome (Fig. 3C) and is reflected also in the key functional enrichments observed for the BAZ1B direct targets that are also expressed and that include “axon guidance”, “tube development”, “dendrite development”, “outflow tract morphogenesis”, “odontogenesis”, “wound-healing” and “endochondral bone morphogenesis” (Fig. 3D). Consistently, the analysis of BAZ1B bound regions uncovered major convergence with the binding motifs of critical NC regulators, including two motifs similar to that of TFAP2A: one *de novo* motif (reverse-engineered from BAZ1B ChIP-seq data and significantly aligning to HOMER motifs database) accounting for 21.49% and one *known* motif (already present in HOMER database and found significantly enriched in BAZ1B ChIP-seq data) accounting for 10.68 % of regions. Among *known* motifs we also found one for NEUROG2, accounting for 14%, and one accounting for 42% of BAZ1B bound regions equally associated to TAL1, TCF12, AP4 and ASCL1 (Fig. 3E). Thus, BAZ1B binding regions are enriched for target sites of major regulators of NC and its neural derivatives (*49*, *50*), among which TFAP2A stands out given its core role in neural borders formation and NC induction and differentiation (*51*) through the binding and stabilization of neural-crest specific enhancers, in concert with NR2F1, NR2F2 and EP300 (*52*).

Finally, we identified 81 regions that are quantitatively bound by BAZ1B depending on its copy-number (FDR < 0.1) (Fig. 3F), 153 regions differentially bound concordantly in WBS and 7dupASD compared to control and atWBS samples (FDR < 0.1) (Fig. S4A), 176 and 25 regions differentially bound preferentially in, respectively, WBS (Fig. S4B) and 7dupASD (FDR < 0.1) (Fig. S4C).

Given the prominence of its binding to distal regulatory regions, we then set out to define the BAZ1B dosage-dependent impact on NCSC-specific enhancers by integrating H3K27ac, H3K4me1, H3K27me3 and H3K4me3 profiles. We thus performed a regression analysis on *BAZ1B* levels for the distribution of the three histone marks in the aforementioned regions and found H3K27ac to be most affected, with 7254 genes differentially acetylated at their enhancers, followed by a differential distribution of the H3K4me1 (4048) and H3K27me3 (2136) marks (Fig. S4D). This enabled the overlay of epigenomic and transcriptomic profiles, uncovering that among the 1192 DEGs identified in the regression RNA-seq analysis, 21.3% (257/1192) are associated to enhancers that are both bound by BAZ1B and differentially H3K27 acetylated in a manner concordant with BAZ1B levels (Fig. S4E), with a stronger overlap for genes whose expression is inversely correlated with BAZ1B levels (160 *vs* 97). The same held for DEGs that have a concordant differential distribution of H3K4me1 mark at enhancers (123 *vs* 55), underscoring the consistency of the impact of BAZ1B dosage on distal regulation (Fig. S4F). In contrast, a lower number of genes (about 30) showed a concordant differential distribution of the H3K27me3 mark and, at the same time, were bound by BAZ1B at enhancers (Fig. S4G), indicating that BAZ1B preferentially impacts active chromatin.

From this integrative analysis, we could thus finally identify a core set of 30 *bona fide* genes that: i) are DEGs in the RNAseq analysis (13 and 17, respectively following BAZ1B levels directly or inversely), ii) have their enhancers differentially marked in a concordant manner (*i.e.*, up in H3K27ac/H3K4me1 and down in H3K27me3 for the upregulated genes, and viceversa for the downregulated) and iii) are bound by BAZ1B at enhancers (Fig. 3G, S4H). This group includes key genes involved in cell migration and adhesion, such as *ARHGEF16*, *NRXN2*, *OLFM1* and *PLXNA4* (*31*, *53*–*55*), as well as cardinal NC markers, such as *HNK1* and *NR2F2* (*56*, *57*). Finally, among the genes whose expression directly follows BAZ1B levels, and whose enhancers are bound by BAZ1B in a quantitatively dosage-dependent manner with a concomitant gain in H3K27ac, we found *PAX5*, which has been implicated in the regulation of neural stem cell proliferation and migration (*58*) and of the developmental balance between osteoblast and osteoclast precursors (*59*). Together, this first dosage-faithful analysis of BAZ1B occupancy in a diverse cohort of human NCSCs establishes its pervasive and mostly distal targeting of the NC-specific epigenome, with a preferentially activator role on the critical transcriptional circuits that define NC fate and function.

### Intersection with paleogenomic datasets uncovers a key evolutionary role for BAZ1B

Mild NC deficits have been put forth as a unifying explanatory framework for the defining features of the so-called domestication syndrome, with BAZ1B listed among the putative underlying genes due to its previously reported role in the NC of model organisms (*3*, *7*, *8*). The recent observation that its expression is impacted by domestication-related mobile element insertion (MEI) methylation in gray wolves (*9*) further supported its role in domestication, offering an intriguing parallel to the paleogenomic results that had detected BAZ1B within the regions of the modern genome reflective of selective sweeps and found it enriched for putatively regulatory mutations in AMHs (*12*). These convergent lines of enticing evidence notwithstanding, neither the neurocristopathic basis of domestication nor its extension to the evolution of the modern human lineage (*i.e.*, self-domestication) have been empirically tested, hence keeping also BAZ1B-based domestication hypotheses to the theoretical ambit.

Having defined the molecular circuits through which BAZ1B shapes the modern human face, we set out to validate the self-domestication hypothesis and its neurocristopathic basis, carrying out a systematic integrative analysis of the overlaps between our empirically defined BAZ1B dosage-sensitive genes (blue Venn in Fig. 4B) and a combination of uniquely informative datasets highlighting differences between modern humans and archaics (Neanderthals/Denisovans) and between wild/domesticated pairs of four paradigmatic species (*1*, *10*–*12*) (represented in Fig. 4A by skulls illustrating the more ‘gracile’ and ‘juvenile’ profile in AMH/dog relative to Neanderthal/wolf visible in the overall shape of the neurocranium, reduced proganthism, browridges and nasal projections). Specifically, as shown in Fig. 4B, these datasets include: i) genes associated with signals of positive selection in the modern branch compared to archaic lineages (purple Venn) (*10*, *11*); ii) genes harboring (nearly) fixed mutations in moderns *vs* archaics (pink Venn); and iii) genes associated with signals of positive selection in the four paradigmatic domesticated species dog, cat, cattle and horse (*1*) (orange Venn), to reveal statistically significant overlaps between them and genes associated with signals of positive selection in the modern branch compared to archaic lineages. In turn, the list of genes harboring (nearly) fixed mutations in moderns *vs* archaics contains three classes: i) genes harboring high frequency changes (*12*) ii) genes harboring missense mutations (red barplot) and iii) genes enriched for mutations in regulatory regions (green barplot) (data based on (*12*)) (Fig. 4C). As shown in the barplots, the obviously very limited number of high-quality coverage archaic genomes available results in a much higher number of nearly fixed changes identified in archaics (left/negative side of the plot) versus modern humans (right side) (Fig. 4C), setting a comparatively much higher threshold for the identification of nearly fixed modern changes.

**Figure 4.**
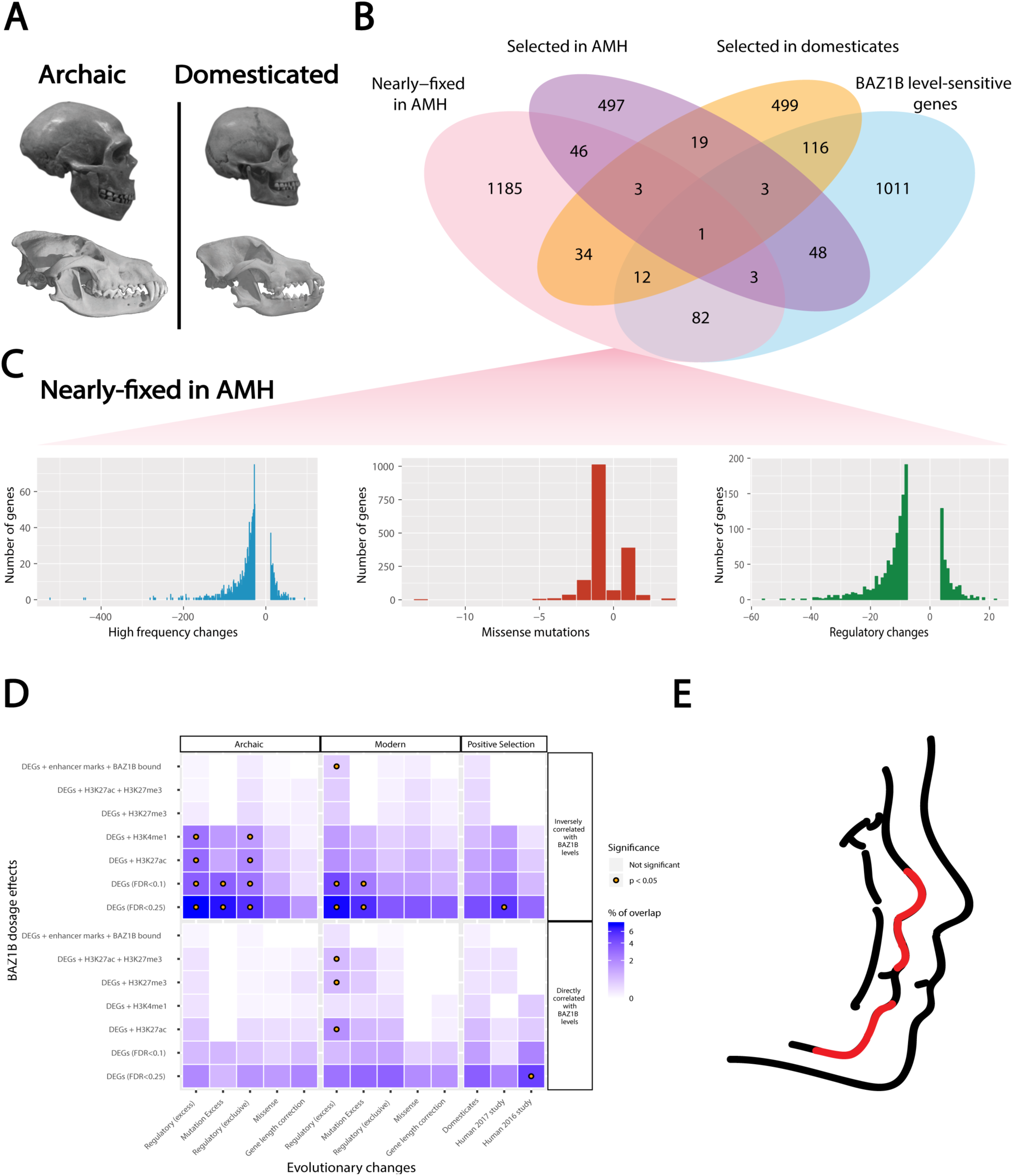
Exploration of paleogenomic datasets supports a key evolutionary role for BAZ1B and validates the self-domestication hypothesis. (**A**) Skulls from archaic human (Neanderthal) and wolf compared to modern human and dog, capturing the parallelism between domestication and self-domestication. (**B**) Overlap between BAZ1B level-sensitive genes and datasets which bring out differences between AMHs and archaics, as well as genes under positive selection in modern humans and domesticates. (**C**) Barplots showing the occurrence of high frequency changes, missense mutations and mutations in regulatory regions in genes from the AMH (nearly) fixed mutation dataset (pink Venn in panel B). (**D**) Heatmap representing the amount of overlaps for each list selected from Panel B. Gene overlaps and detailed list descriptions are reported in Tables S1-S14. (**E**) Rendering of a typical WBS face (left) against the background of a typical modern face (right). Red segments indicate areas of the lower face where the two faces most sharply depart (nose, philtrum, lower front of the mandible). The lower mid-face region is the most often associated with mutations in genes figuring prominently in our intersections, as discussed in the text and Table S15.

These analyses are visualized in Fig. 4D (and detailed in Tables S1-15) through a matrix that intersects all BAZ1B dosage-dependent genes (partitioned in the two categories of directly and inversely correlated, and ordered across the full range of biological significance and regulatory proximity, from simply differentially expressed genes to bona fide direct targets) with the evolutionary changes underlying domestication and self-domestication, yielding the following key insights (color coded for degree of overlap and marked for significance in hypergeometric tests). First, the most significant pattern obtains at the intersection with the top 10% genes showing an excess of (nearly) fixed mutations in the regulatory regions of modern humans compared to archaics, across both directly and inversely BAZ1B level-dependent genes (Tables S1-14). Importantly, this same category of nearly fixed modern regulatory changes is also the only one that returns a statistically significant overlap with the most stringent core of BAZ1B dosage dependent targets (ie. DEGs whose enhancers are both directly bound by BAZ1B and differentially marked upon its decrease), demonstrating that BAZ1B directly controls, in an exquisitely dosage-dependent manner, this particularly relevant set of genes that underwent regulatory changes in human evolution. Second, the overall strongest overlaps map to the class of genes that are inversely correlated to BAZB levels, which we found to be strongly and specifically enriched for head morphogenesis and NC categories (Fig. 2C), thereby confirming craniofacial morphogenesis as the key domain of functionally relevant overlap between BAZ1B dosage and (self-)domestication changes relevant to the evolution of AMHs. Third, despite the spuriously inflated number of apparently fixed mutations in archaics (*12*), the overall extent of overlap between genes impacted by BAZ1B dosage and our modern and archaic sets does not reveal significantly more hits for archaics. In fact, globally we found consistently more overlapping genes between the BAZ1B targets and the modern human data, and even no statistically significant overlap for any list of the archaic-specific mutations when crossed against BAZ1B level-directly correlated genes. Finally, the (lower) midface emerges as a particularly salient area of functionally relevant overlap (as illustrated in Fig. 4E and detailed in Table S15), given the specific genes that our analysis unearthed: i) *COL11A1*, one of the few craniofacial genes highlighted across domestication studies (dog, house sparrow, pig breed) and which lies in a region of the human genome that resisted archaic introgression (*10*), is associated with Marshall syndrome and was also highlighted in a recent study on regulatory changes that shaped the modern human facial and vocal anatomy (*60*); ii) *XYLT1*, one of the 5 genes (along with *ACAN*, *SOX9*, *COL2A1*, and *NFIX*) that affect lower and midfacial protrusion and are among the top differentially methylated genes compared to archaics (Table S15); and iii) *NFIB*, which belongs to the same gene family as *NFIX* and shares some of its functions.

## Discussion

Following on his seminal account of domestication systematized in *Variations of Animals and Plants under Domestication* (*61*), Charles Darwin first put forward the analogy between modern humans and domesticated species in *The Descent of Man* (*62*), although his emphasis on controlled breeding as a key aspect of domestication left his intuition largely unresolved (*63*). Since then the possibility that the anatomical and cognitive-behavioral hallmarks of AMHs could result from an evolutionary process bearing significant similarities to the domestication of animals has been refined into the full-fledged self-domestication hypothesis (*1*, *2*), spurring considerable interest but failing to achieve epistemic closure largely due to two factors: i) the lack of a coherent explanation, even at a theoretical level, of what developmental and genetic mechanisms could underlie domestication in general; and ii) the absence of suitable experimental systems in which those mechanisms could be specifically tested in the case of human self-domestication. The first problem was tackled by the recent proposition of mild NC deficits as a central and unifying functional layer underlying domestication (*3*). This constituted a major conceptual advance, not least because it generated for the first time the testable hypothesis of an altered NC gene expression program in domesticated species relative to their wild-type ancestors. For humans, given the obvious lack of gene expression data from archaic hominins, we reasoned that this hypothesis could be verified by interrogating the genetic changes between archaic and modern humans with the gene regulatory networks directly inferred from human neurocristopathies. We thus set out to test whether specific human neurodevelopmental disorders, carefully selected on the basis of both craniofacial and cognitive-behavioral traits relevant to domestication, could illuminate the regulatory circuits shaping the modern human face and hence be harnessed for an empirical validation of the self-domestication hypothesis. Specifically, we reasoned that WBS and 7dupASD, through their uniquely informative set of symmetrically opposite phenotypes at the level of face morphology (Fig. S1A) and sociality, constituted a paradigmatic test case to probe the heuristic potential of neurodevelopmental disease modelling for the experimental understanding of human evolution. The following key insights confirm the validity of this approach.

First, we identified the 7q11.23 region *BAZ1B* gene as a master regulator of the modern human face on the basis of a molecular and functional dissection in the thus far largest cohort of WBS and 7dupASD patient-specific NCSCs and across an exhaustive range of BAZ1B dosages. Notably, our cohort also included NCSCs from a rare WBS patient featuring a much milder WBS *gestalt* and harboring an atypical, BAZ1B-sparing deletion that served as a particularly informative control, as confirmed by the clustering of atypical NCSC lines with controls when probed for BAZ1B occupancy. In particular, exploiting the fine-grained resolution of BAZ1B dosages recapitulated in our cohort, we could couple classical pairwise comparisons with a more sophisticated regression analysis on BAZ1B levels, thereby revealing major BAZ1B dosage-dependent transcriptional alterations pivoting around clusters of pathways that are crucial for NC development and maintenance, as well as for its downstream skeletal and cardiac outputs. Second, we repurposed the versatility of CRISPR/Cas9 to generate an allelic series of endogenously tagged BAZ1B across 7q11.23 dosages (including the BAZ1B-sparing atypical patient as uniquely relevant control) to define its dosage-dependent genome-wide occupancy. Taking advantage of previous extensive work on the NCSC chromatin landscape (*36*, *52*, *56*, *64*), we were able to define a pivotal role for BAZ1B in NCSC enhancer regulation, consistent with its preferential binding of distal regulatory regions, and to partition its dosage-dependent regulation into *bona fide* direct and indirect targets. Interestingly, the overall balance between the numbers of genes up-or down-regulated upon BAZ1B KD, together with the greater overlap, sheer size and significance of enrichments in chromatin remodeling categories over other domains of transcription regulation, further corroborates the inclusion of BAZ1B among the factors acting upstream of enhancer and promoter modulations to enable or reinforce rather than specify their net outcome. Finally, this molecular readout was translated to the functional level with the definition of an impairment in both NCSC migration and outgrowth from EBs upon decrease in BAZ1B, providing the first validation of BAZ1B involvement in key functions of the developing human NCSCs.

Third, our investigation provides the first experimental evidence for the neurocristopathic hypothesis that had been put forth to explain domestication and had pointed to BAZ1B as one of the candidates underlying the domestication syndrome (*3*). Indeed, among the key NC hubs impacted by BAZ1B dosage, we uncovered three additional critical genes, *EDN3*, *MAGOH*, and *ZEB2*, that had also been predicted in the same model because they are associated with behavioral changes found in domesticates, thereby defining a regulatory hierarchy for this coherent set of genes underlying domestication.

Finally, the empirical determination of BAZ1B dosage-sensitive genes, in NC models from AMHs with accentuated domestication-relevant traits, allowed us to functionally interrogate datasets exposing the genetic differences between modern *vs* archaic. This brought into relief the significant convergence between BAZ1B-dependent circuits and genes harboring regulatory changes in the human lineage, reinforcing the notion that regulatory regions contain some of the most significant changes relevant for the modern lineage. In this context, it is noteworthy that genes implicated in NC development also play significant roles in the establishment of brain circuits that are critical for cognitive processes like language or theory of mind prominently affected in 7q11.23 syndromes. Indeed, among the genes downstream of BAZ1B that we uncovered in this study, *FOXP2*, *ROBO1* and *ROBO2*, have long been implicated in brain wiring processes critical for vocal learning in several species (*65*, *66*), including humans, and will warrant further mechanistic dissection in light of the distinctive linguistic profile of WBS individuals.

Thus, our findings establish the heuristic power of neurodevelopmental disease modelling for the study of human evolution.

## Acknowledgements

We thank the imaging and genomic units at IEO and IFOM for the help with wound-healing assays and sequencing, respectively, and Prof. Giorgio Scita and his group for suggestions on migration experiments. **Funding**: This work was funded by the Telethon Foundation (grant number GGP14265 to G.T. and G.M.), the EPIGEN Flagship Project of the Italian National Research Council (to G.T.), the European Research Council (consolidator grant number 616441-DISEASEAVATARS to G.T.), the Horizon 2020 Innovative Training Network EpiSyStem, Ricerca Corrente granted by the Italian Ministry of Health (G.T. and G.M.), Daunia Plast (to G.M.), Fondazione Umberto Veronesi (to P.L.G.), the IEO Foundation (fellowship to A.V.), the Spanish Ministry of Economy and Competitiveness/FEDER funds (grant FFI2016-78034-C2-1-P to C.B.), the Generalitat de Catalunya (grant 2017-SGR-341 to C.B. and doctoral fellowship to T.O.), the MEXT/JSPS Grant-in-Aid for Scientific Research on Innovative Areas 4903 (Evolinguistics: JP17H06379 to C.B.), a Marie Curie International Reintegration Grant from the European Union (PIRG-GA-2009-256413 to C.B.), the European Social Fund (grant BES-2017-080366 to A.A.) and the Portuguese Foundation for Science and Technology (PhD grant number SFRH/BD/131640/2017 to P.T.M.). **Author contributions:** M.Z. carried out all the experiments including the generation and characterization of both BAZ1B-interfered lines and BAZ1B-tagged lines, the preparation of RNA-seq libraries, ChIPs and wound-healing assays; A.V. performed RNA-seq and ChIP-seq bioinformatic analyses with the supervision of P.-L.G.; A.A., P.T.M., S.S., T.O. and C.B. performed analyses in Fig. 4; M.L. and A.R.-I. characterized NC induction phenotype reported in Fig. 1D,E; N.M. reprogrammed 7dupASD3 and CTL4R fibroblasts and generated the corresponding NC lines; A.S. helped with cell culture; M.Z. and S.T. designed the tag strategy; G.M. provided 7dupASD3 and CTL4R fibroblasts; M.Z., C.B. and G.T. wrote the manuscript with contributions from A.V. and A.S.; G.T. conceived, designed and supervised the study. **Competing interests**: None. **Data and materials availability**: All data are available in the manuscript or the supplementary materials.

## Supplementary Materials

Materials and Methods

Figures S1-S4

Tables S1-S17

References (67-85)

