## Supplementary materials and figures for "7q11.23 Syndromes Reveal BAZ1B as a Master Regulator of the Modern Human Face and Validate the Self-Domestication Hypothesis"

**This PDF file includes:**

Materials and Methods

Figs. S1 to S4

Tables S1 to S17

#### Material and Methods

Human samples

Ethics approvals are reported in the study that established the original iPSC cohort (*13*) and also apply to the additional samples included in this study (7dupASD3 and CTL4R).

Fibroblast reprogramming and iPSC culture

WBS1, WBS2, WBS3, WBS4, 7dupASD1, AtWBS1 and CTL2 fibroblasts were reprogrammed using the mRNA Reprogramming kit (Stemgent), while the 7dupASD2 and CTL1R lines were reprogrammed with the microRNA Booster kit (Stemgent). The CTL3 line was reprogrammed by transfection with the STEMCCA polycistronic lentiviral vector followed by Cre-mediated excision of the integrated polycistron. 7dupASD3 and CTL4R fibroblasts were reprogrammed using the Simplicon$^{TM}$ RNA Reprogramming kit (Millipore).

Prior to differentiation, iPSC lines were cultured on hESC-qualified Matrigel (BD Biosciences) coated plates, diluted 1:40 in DMEM/F-12 and grown in mTeSR-1 medium (StemCell Technologies). They were passaged upon treatment with Accutase (Sigma) and then plated in mTeSR-1 medium supplemented with 5µM Y-27632 (Sigma).

Differentiation

Differentiation into NCSCs was performed as previously described (*67*), with the exception of NCSCs used in the experiment reported in Fig. 1D-F (*16*).

Flow cytometry

NCSCs were detached using accutase, counted and $1\times10^6$ cells/experimental condition were fixed in 4% paraformaldehyde and then blocked in 10% Bovine Serum Albumin (BSA). Cells were incubated for 1 hour with primary antibodies conjugated to fluorophores (HNK1-FITC and NGFR-ALEXA647,

BD Biosciences). Analyses were performed on a FACSCalibur instrument (BD Biosciences) and data were analyzed with FCS express software (Tree Star).

FACS characterization for 7dupASD3 and CTL4R lines is reported in Fig. S1B; for all the other lines see (*13*).

Lentiviral vector production and NCSC transfection

*BAZ1B* knock-down was performed using validated pLKO.1 TRC vector TRCN0000013338 (referred to as sh1) and TRCN0000013341 (referred to as sh2). A pLKO.1 TRC vector containing a scrambled short hairpin sequence was used as a negative control.

Second generation lentiviral vectors were produced through calcium phosphate transfection of human embryonic kidney (HEK293T) cells and ultracentrifugation (2 hours, 20°C, 20000 rpm).

3-4x$10^5$ NCSCs were infected upon splitting and then selected by adding 1µg/ml puromycin to the medium.

RNA extraction, retrotranscription and Real Time quantitative PCR (RT-qPCR)

RNA was extracted using the RNeasy Micro Plus kit (Qiagen) according to manufacturer's instructions.

Retrotranscribed cDNA was obtained from 0.5-1 µg of total RNA using the SuperScript VILO Retrotranscription kit (Thermo Fisher Scientific).

RT-qPCR was performed on a 7500 Fast Real-Time PCR system (Applied Biosystems) using SYBR Green Master Mix (Applied Biosystems) as detecting reagent. A total cDNA amount corresponding to 15 ng of starting RNA was used for each reaction. Each sample was analyzed in triplicate and normalized to *GAPDH*. Relative mRNA quantity was calculated by the comparative cycle threshold (Ct) method using the formula $2^{-deltaCt}$.

*BAZ1B*: CCTCGCAGTAAGAAAGCAAAC (forward); ACTCATCCAGCTCCTTTTGAC (reverse).

*GAPDH*: GCACCGTCAAGGCTGAGAAC (forward); AGGGATCTCGCTCCTGGAA (reverse).
*NR2F1*: AGAAGCTCAAGGCGCTACAC (forward); GGGTACTGGCTCCTCACGTA (reverse).
*NR2F2*: GCAAGTGGAGAAGCTCAAGG (forward); GCTTTCCACATGGGCTACAT (reverse).
*TFAP2A*: GCCTCTCGCTCCTCAGCTCC (forward); CGTTGGCAGCTTTACGTCTCCC (reverse).
*SOX9*: AGTACCCGCACTTGCACAAC (forward); GTAATCCGGGTGGTCCTTCT (reverse).

RNA-seq libraries preparation

Library preparation for RNA sequencing was performed according to TruSeq Total RNA sample preparation protocol (Illumina), starting from 250 ng - 1 µg of total RNA.
cDNA library quality was assessed at an Agilent 2100 Bioanalyzer, using the High sensitivity DNA kit.
Libraries were sequenced with the Illumina HiSeq machine at a read length of 50 bp paired-end and a coverage of 35 million of reads per sample.

Protein extraction and Western Blot

NCSCs were lysed in RIPA buffer (10 mM Tris, pH 8.0, 1% Triton X-100, 0.1% sodium deoxycholate, 0.1% SDS, 140 mM NaCl, 1 mM EDTA) supplemented with protease inhibitor cocktail (Sigma) and 0.5 Mm Phenylmethylsulfonyl fluoride (PMSF) (Sigma) for 1 hour at 4°C.
Protein extracts (30-50 µg per sample) were supplemented with NuPAGE LDS sample buffer (Thermo Fisher Scientific) and 50 mM Dithiothreitol (DTT) (Thermo Fisher Scientific) and denaturated at 95°C for 3 minutes. Then extracts were run on a precast NuPAGE 4–12% Bis-Tris Gel (Thermo Fisher Scientific) in NuPAGE MOPS SDS Running Buffer (Thermo Fisher Scientific) and transferred to a 0.45 µm nitrocellulose membrane (GE Healthcare) for one hour at 100 V in a buffer containing 20% absolute ethanol and 10% 0.25 M Tris base, 1.9 M glycine. The membranes were blocked in TBST (50 mM Tris, pH 7.5, 150 mM NaCl and 0.1% Tween- 20) and 5% milk for one

hour, incubated with primary antibodies overnight (O/N) at 4°C and with secondary antibodies for one hour at room temperature (RT). Primary (BAZ1B, Abcam and GAPDH, Millipore) and secondary antibodies were diluted in TBST and 5% milk. Blots were detected with the ECL Prime Western Blotting detection reagents (Sigma) and scanned using ChemiDoc system (Biorad).

Wound-healing assay

5-7x$10^4$ cells were plated in each of the two matrigel-coated wells of silicone culture-inserts (Ibidi) attached to 6 well-culture plates. After 24 hours, the insert was removed, medium was changed to remove dead cells and time lapse was performed for 24 hours at the rate of one image every 10 minutes at a 10x magnification; each condition was analyzed in duplicate. Images were acquired with the BX61 upright microscope equipped with a motorized stage from Olympus or the Nikon Eclipse Ti inverted microscope equipped with a motorized stage from Nikon and analyzed with ImageJ.

Endogenous BAZ1B tagging via CRISPR/Cas9

iPSCs were pretreated with 10 µM Rock inhibitor for 4 hours and then 2x$10^6$ cells were electroporated using the NEON system with the Cas9/sgRNA ribonucleoprotein complex and the donor plasmid (synthesized by GeneArt). The donor plasmid contains three Flag tag followed by a self-cleaving peptide (P2A) and a Hygromycin resistance (HygroR). The 3xFlag-P2A-HygroR cassette was flanked by BAZ1B specific Homology Arms (5' HA and 3' HA) to promote homologous recombination and then subcloned into a bacterial backbone (Fig. 3A).

After 48 hours iPSC medium was supplemented with 50 µg/µl Hygromycin B and selection medium was maintained for 15 days. 15-20 clones per iPSC line where then subjected to PCR to i) evaluate the presence of the cassette and the insertion in the correct genomic locus and ii) distinguish heterozygously from homozygously tagged clones (Fig. S3A). We could isolate a clone with a homozygous integration from the CTL, the atypical WBS and the typical WBS, but not from the

7dupASD line. In the 7dupASD clone the Flag tag was present in 2 out of 3 copies, as shown by a digital PCR analysis (Fig. S3B).

Digital PCR

60 ng of DNA was amplified in a reaction volume containing the following reagents: "QuantStudio™ 3D Digital PCR Master Mix v2 (Thermofisher), "Custom TaqMan Copy Number Assays, SM 20x, FAM labeled (Thermofisher) and "TaqMan Copy Number Reference Assay 20x (Thermofisher) VIC labeled (Thermofisher) . The mix was loaded on a chip using the QuantStudio 3D Digital PCR Chip Loader. The chips were then loaded on the Proflex PCR System (Thermofisher) and data were analyzed using the "QuantStudio 3D Analysis Suite Cloud Software". The entire process was performed by the qPCR-Service at Cogentech-Milano.

Custom (FLAG) TaqMan Copy Number Assays:

FW Primer TGGACAGTCCAGAGGACGAA

RV Primer CACCCTTGTCGTCATCGTCTT

Probe FAMACAGAAGAAGGACTACAAAGACG

TaqMan Copy Number Reference Assay:

TERT (VIC) (catalog number 4403316)

Chromatin immunoprecipitation coupled with sequencing (ChIP-seq)

Approximately 2 x $10^5$ cells were used (~100 ug of chromatin) for histone mark immunoprecipitation and 1 mg of chromatin for BAZ1B-FLAG immunoprecipitation. Cells were fixed with PBS, containing 1% formaldehyde (Sigma), for 10 minutes to cross-link proteins and DNA, when the reaction was then stopped by adding 125 mM glycine for 5 minutes. Cells were lysed with SDS buffer containing 100 mM NaCl, 50 mM Tris-HCl pH 8.0, 5 mM EDTA pH 8.0, 10% SDS, at which point chromatin pellets were resuspended in IP buffer containing 1 volume of SDS buffer and 0.5 volume of Triton dilution buffer (100 mM Tris-HCl pH 8.5, 5 mM EDTA pH 8.0, 5% Triton X-100). Chromatin was then sonicated using the S220 Focused-ultrasonicator (Covaris) to generate <300 bp DNA fragments (for histone mark IPs) or the Branson digital sonifier to generate 500-800 bp DNA fragments (for BAZ1B-FLAG IP).

Sonicated chromatin was incubated O/N at 4°C with primary antibodies (H3K27ac, Abcam; H3K4me1, Abcam; H3K4me3, Abcam; H3K27me3, Cell Signaling and FLAG, Sigma) and then 3 hours with Dynabeads Protein G (Thermo Fisher Scientific). Beads were washed three times with low-salt wash buffer (0.1% SDS, 1% Triton X-100, 2 mM EDTA, 20 mM Tris-HCl pH 8.0 and 150 mM NaCl) and once with high-salt wash buffer (0.1% SDS, 1% Triton X-100, 2 mM EDTA, 20 mM Tris-HCl pH 8.0 and 500 mM NaCl). Immunocomplexes were eluted in decrosslinking buffer (1% SDS, 100 mM NaHCO3) at 65 °C for 2 hours, DNA was purified using QIAquick PCR columns (Qiagen) and quantified with Qubit dsDNA HS assay kit (Thermo Fisher Scientific). DNA libraries were prepared by the sequencing facility at IEO campus according to the protocol described by Blecher-Gonen and colleagues (*68*) and DNA was sequenced on the Illumina HiSeq 2000 platform. For the FLAG ChIP, samples were run in duplicate.

RNA-seq analysis

RNA-seq data were quantified using Salmon 0.91 to calculate read-counts and transcripts per million (TPMs) in a transcript- and gene-wise fashion, using the quasi-mapping off-line algorithm (*69*) on GRCh38 (NCBI) database. EdgeR was used for differential gene expression analysis (DEA), using generalized linear regression methods (GLMRT), to identify pattern of differential expression following two different schemes:

1. a factorial analysis based on the definition of one group of scrambled and one group of knock-down samples to identify genes dysregulated similarly across short-hairpins characterized by different efficiencies;

2. a numerical analysis in which log-normalized (TMM) BAZ1B levels, as quantified by RNA- seq, were used as independent variable.

All analyses were performed dropping individual variations (~individual+KD or ~individual+BAZ1B), in order to account for the genetic background of each individual. In particular, this design is expected to permit the identification of genes which change expression level upon KD even in situations in which genotype- specific make-ups would lead BAZ1B-dependent genes to have unique expression levels in scramble lines. In the factorial analysis, differentially expressed genes (DEGs) were identified and characterized by filtering for fold-change (FC) > 1.25 and FDR < 0.05 unless explicitly indicated.

To our knowledge, performing a regression analysis at a gene-specific level has never been performed. We were able to do this because of the availability of a large set of samples (11 individuals) and because of the two short-hairpins robustly respectively reducing BAZ1B expression levels, respectively by ~40% and ~70% in all individuals lines. To validate the quality of our numerical differential expression analysis we took advantage of HipSci data (*70*, *71*) and iPSCpoweR tools (*26*). We took 50 out of 105 possible combinations of 13 random individuals RNA-seq data from the healthy HipSci cohort, representing both sexes and having at least two technical replicates per individual. Unfortunately, HipSci does not contain at least 13 individuals with 3 clones per individual. Thus, we performed four alternative DEAs with edgeR (Table S16) on the 50 different random combinations of 13 individuals identified (200 DEAs in total, on 22 samples, 2 clones per individual), using the same model matrix used for the regression analysis (~individual+BAZ1B), using BAZ1B levels of scramble and sh2 lines. All analyses identified very low number of spurious DEGs (Fig. S2E). Thus, we used the “Edg2” pipeline (Table S16), because it does not discard genes with higher variability (Edg2 and Edg4 vs Edg1 and Edg3) and it is based on a better suited algorithm (Edg2 vs Edg4). With our model matrix, filtering by p-value < 0.01 (and FDR < 0.25), using Edg2 on a random HipSci data, we obtained an average of 93.32 differentially expressed genes (on average) with a median equal to 43 (Table S17).

Gene Ontology (GO) enrichments were performed using topGO R package version 2.28.0.

Master regulatory analysis was performed via hypergeometric test, by measuring geneset enrichments in lists of transcription factor (TF) targets provided by TFBS tools database (*72*). Both GO and TF enrichment analyses were performed considering as background genes expressed in at least 2 samples in our NCSC cohort.

ChIP-seq analysis

ChIP-seq experiments were analyzed both qualitatively and quantitatively. Reads were trimmed with the FastX toolkit (-Q33 -t 20 -l 22), aligned with Bowtie 1.0 (-v 2 -m 1) on the Human hg38 reference genome and peaks were called by means of MACS 2.1.1. H3K4me1, H3K27ac, H3K4me3 and H3K27me3 peaks were called with --broad using default parameters and $q < 0.05$.

Qualitative analysis, including intersection and comparison of bed files, was performed using BedTools version 2.23.

To define enhancer regions, we intersected those marked by H3K4me1 and H3K27ac in at least two samples, discarded regions with H3K4me3 in at least two samples, and discarded regions overlapping with TSS.

Motif enrichment was performed by means of Homer v4.10.

Quantification of reads per region was performed with DeepTools 3.0.2. Differential mark deposition was conducted by means of edgeR 3.24.1 inside R 3.3.3. To define mark deposition following BAZ1B levels we used the same design as for RNA-seq data (~individual+BAZ1B).

In order to identify BAZ1B bound regions, and to avoid losing identification of lowly covered regions, we resorted to i) aggregation of all samples aligned reads and ii) peak calling with MACS2 using –extsize 800 and $q < 0.25$. BAZ1B binding coverage was calculated with DeepTools, with the same parameters used for histone marks, on the identified peak regions. Differentially bound regions were identified with edgeR.

Assembly of archaic and modern human lists

The archaic/modern lists were generated from the material presented in (*12*). We used high-coverage genotypes for three archaic individuals: one Denisovan (*73*), one Neanderthal from the Denisova cave in Altai mountains (*74*) and another Neanderthal from Vindija cave, Croatia (*75*). The data is publicly available at http://cdna.eva.mpg.de/neandertal/Vindija/VCF/, with the human genome version hg19 as reference. High frequency differences were defined as positions where more than 90% of present-day humans carry a derived allele, while at least the Denisovan and one Neanderthal carry the ancestral allele. High-frequency changes in archaics were defined as occurring at less than 1% in present-day humans, while at least two archaic individuals carry the derived allele. The HF lists used here were examined as presented in (*12*), with the exception of the HF lists in regulatory regions which were extracted from the same dataset, but not presented as such in the original paper.

### Supplementary Figure 1.

**A**

| WBS | 7dupASD | |
|---|---|---|
| Low nasal root | High, broadnose | Symmetrically opposite |
| Long philtrum | Short philtrum | Symmetrically opposite |
| Full lips | Thin lips | Symmetrically opposite |
| Broad forehead | | Shared |
| Dental abnormalities | | Shared |

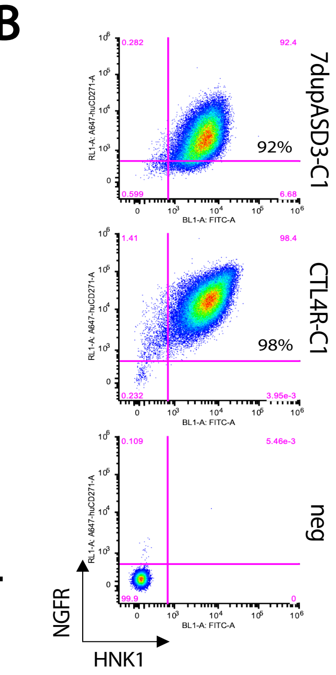


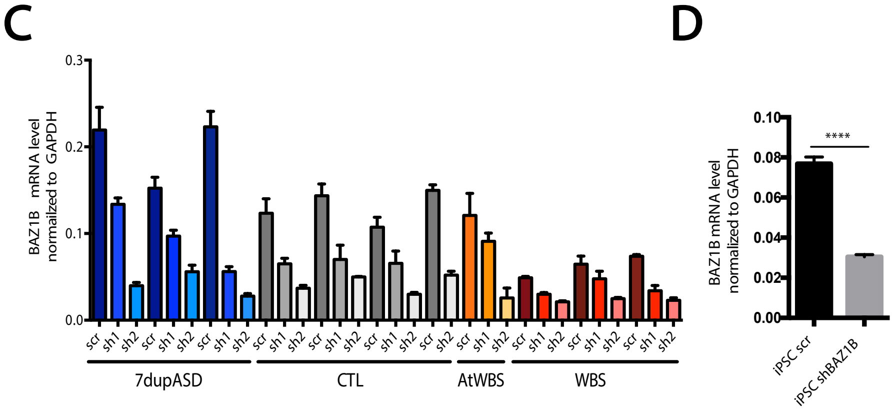


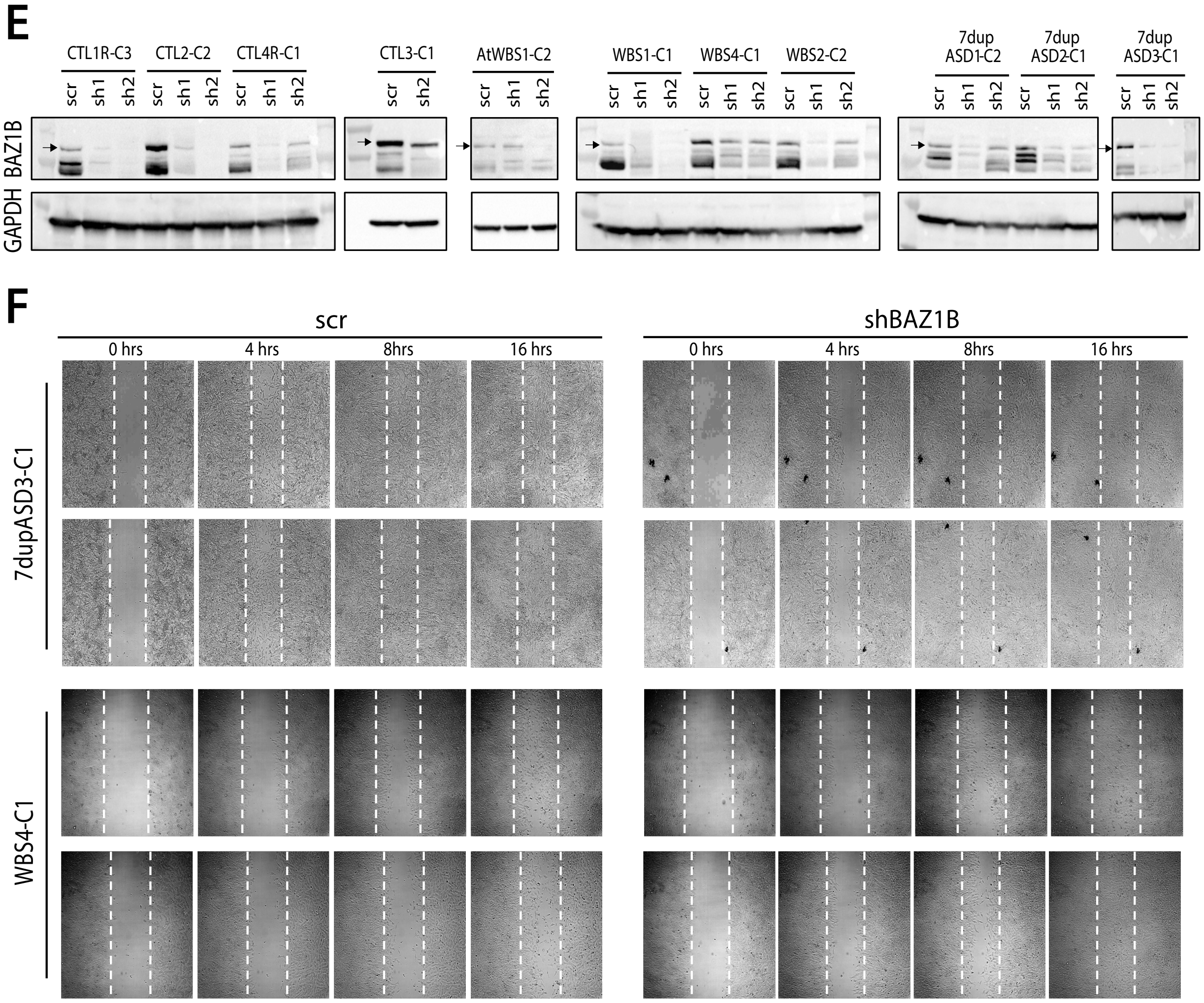

**Figure S1. Generation and BAZ1B KD validation in iPSC-derived NCSCs. (A)** Symmetrically opposite and shared craniofacial phenotypes in WBS and 7dupASD patients**. (B)** Flow cytometry analysis of $HNK1^{+}/NGFR^{+}$ NCSCs (7dupASD3-C1 and CTL4R-C1). Unstained cells are used as negative control**. (C)** BAZ1B mRNA levels in all the interfered lines (scr, sh1 and sh2) as measured by qPCR. Data represent individual samples with the same number of BAZ1B copies. GAPDH is used as normalizer. **(D)** qPCR validation of BAZ1B KD in the iPSC line used in the experiment reported in Fig. 1D, E. **(E)** Western blots showing BAZ1B levels upon KD in all NCSC lines. The arrow indicates the BAZ1B specific band. GAPDH is used as the normalizer. **(F)** 4 hours, 8 hours and 16 hours-time points from the wound-healing assay analysis performed on a 7dupASD and a WBS NCSC line upon BAZ1B KD. Cells from the same line infected with the scr sh were used as references for the migration (n=2).

### Supplementary Figure 2.

A

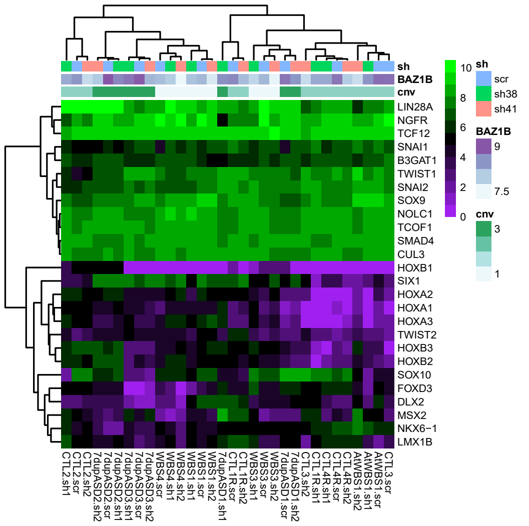


B

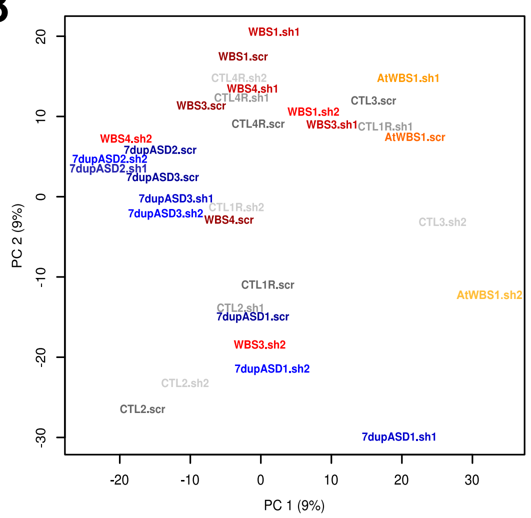


C

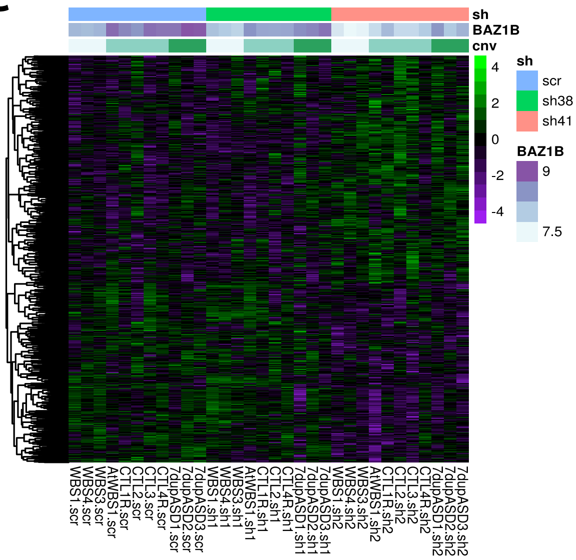


D

| GENE | % of BAZ1B DEGs | GENE | % of BAZ1B DEGs |
|---|---|---|---|
| CEBPB | 52,46 | TBP | 66,74 |
| RAD21 | 86,16 | TAF1 | 73,44 |
| CTBP2 | 31,25 | MAX | 41,07 |
| USF1 | 66,96 | SP1 | 52,00 |
| TEAD4 | 54,69 | EGR1 | 37,50 |
| RBBP5 | 64,73 | KDM1A | 27,46 |
| MAFK | 37,50 | SIN3A | 70,09 |
| SRF | 28,79 | MXI1 | 31,03 |
| CTCF | 82,14 | USF2 | 27,68 |
| TCF12 | 32,10 | CHD2 | 31,47 |
| YY1 | 62,05 | POLR2A | 69,42 |
| HDAC2 | 26,79 | SP4 | 25,22 |
| EP300 | 37,50 | TAF7 | 40,63 |
| GTF2I | 31,25 | | |

E

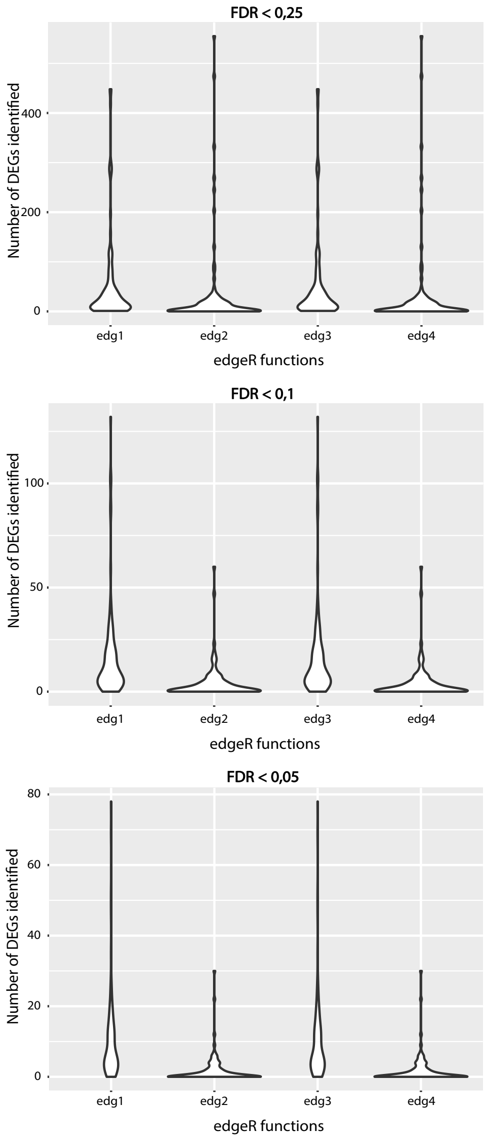

**Figure S2. BAZ1B knock down affect the transcriptome of iPSC-derived NCSCs. (A)** Expression levels (log-TMM) of cranial- and NC-specific genes constituting the signature used to assess the quality of our NCSC lines. (**B**) Principal component analysis showing the distribution of the 32 NCSC lines according to their transcriptional profiles. **(C)** Expression profile of the 448 genes that follows BAZ1B levels. The samples are ordered according to shRNAs. In each shRNA group samples are further ordered based on *BAZ1B* copies (CNVs) (FDR < 0.1). **(D)** List of genes identified as regulators of BAZ1B levels-sensitive genes in the Master regulator analysis. For each gene the percentage of BAZ1B-regulated DEGs is reported. (**E**) Violin Plot representation of the number of spurious DEGs generated by the four implemented edgeR pipelines on HipSci RNA-seq data, given the model matrix applied for the regression on BAZ1B levels in NCSCs.

### Supplementary Figure 3.

A

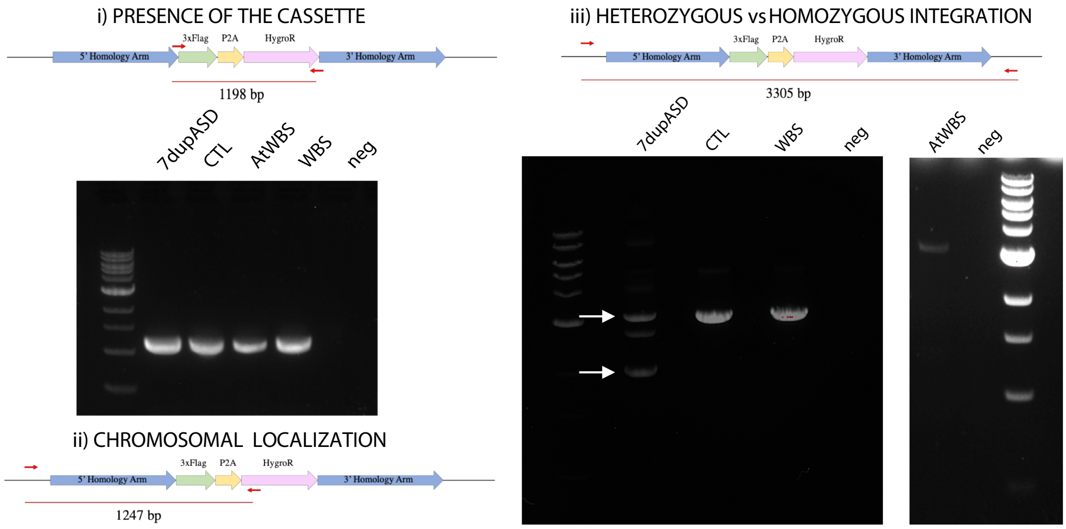


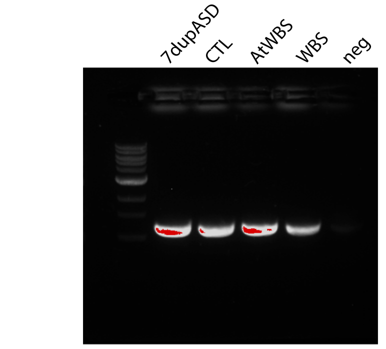


B

| SAMPLE | Copies/µl (VIC) | Copies/µl (FAM) | FLAG copies |
|---|---|---|---|
| 7dupASD | 2576,1 | 2771,4 | 2,1516 |
| CTL | 139,23 | 150,14 | 2,1567 |
| atWBS | 574,42 | 570,57 | 1,9865 |
| WBS | 372,45 | 221,11 | 1,1873 |
| neg | 19,218 | 1,591 | |

C

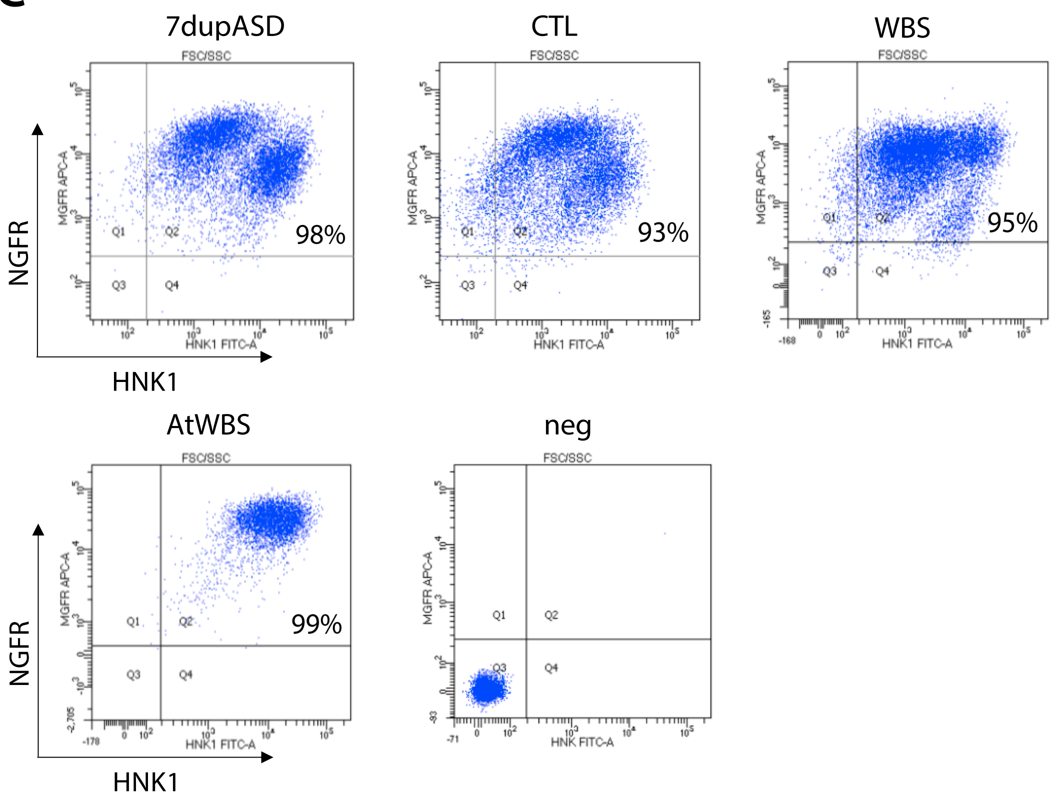

**Figure S3. Generation and differentiation to NCSCs of BAZ1B-FLAG iPSC lines.** (**A**) PCR reporting FLAG-tagged selected clones. Clones were screened for i) the presence of the cassette (band at 1198 bp) and ii) its proper chromosomal localization (band at 1247 bp) and iii) to distinguish a heterozygous (2 bands: one at 2138 bp and one at 3305 bp) from a homozygous integration (single band at 3305 bp). (**B**) Digital PCR analysis to evaluate the exact number of FLAG integrated copies. The FLAG probe was labeled with FAM, while a TERT probe labeled with VIC was used as an internal reference (two known copies). (**C**) FACS analysis of HNK1$^{+}$/NGFR$^{+}$ iPSC-derived NCSC tagged clones. Unstained cells were used as negative control.

### Supplementary Figure 4.

A B C

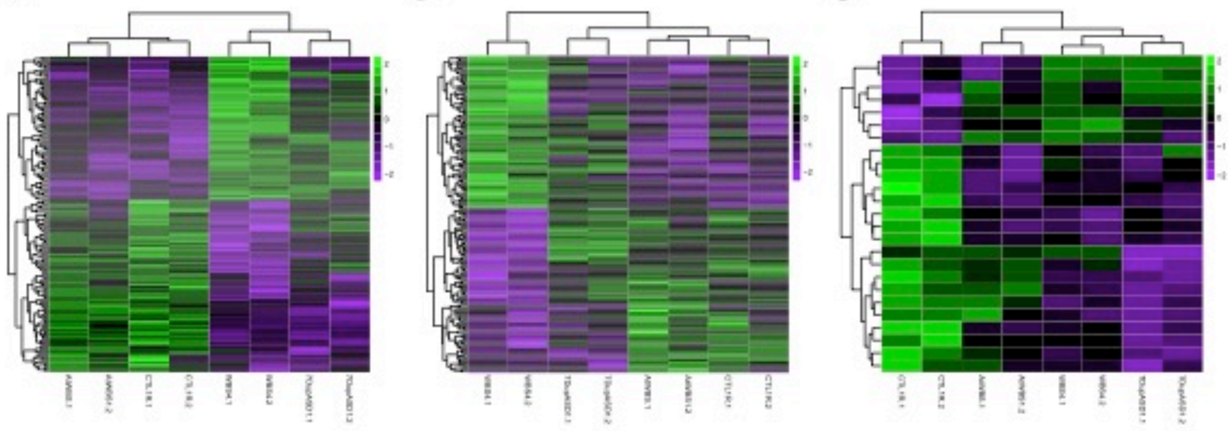

D

| HISTONE MARK | DIRECT | INVERSE |
|---|---|---|
| H3K27ac | 3667 | 3587 |
| H3K4me1 | 1879 | 2169 |
| H3K27me3 | 1039 | 1097 |

E

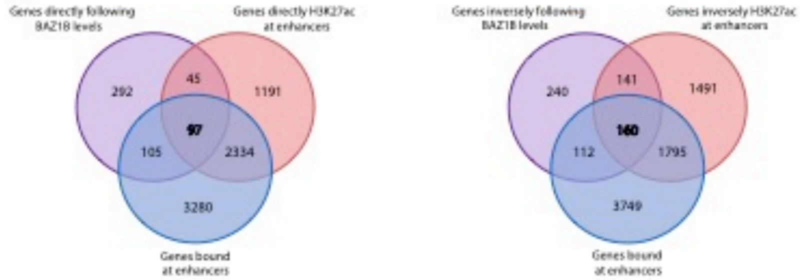


F

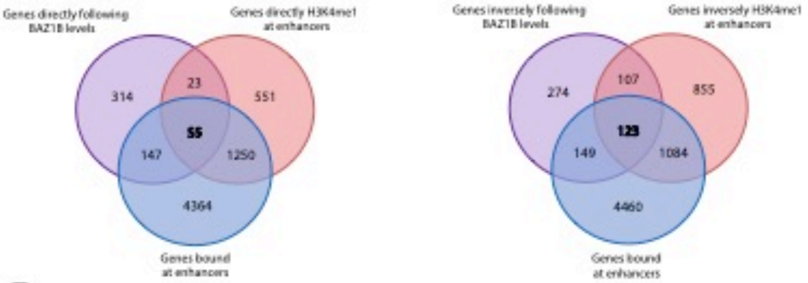


G

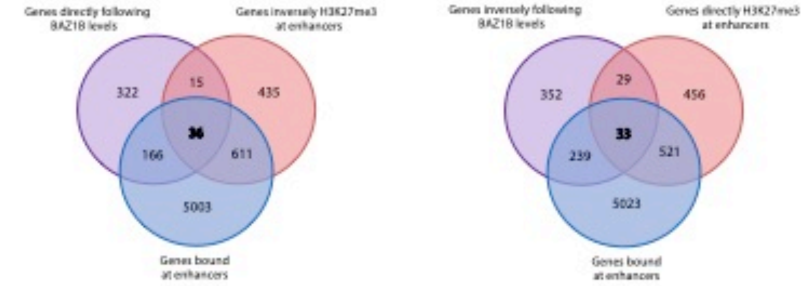


H

| DIRECT | INVERSE |
|---|---|
| ABLIM2 | ADORA2B |
| ARHGEF16 | CHST15 |
| B3GAT1 | COL12A1 |
| DOK7 | DOCK9 |
| IL4R | GMPR |
| KHDRBS3 | KLF7 |
| MGAT5B | MAFB |
| NRXN2 | MRC2 |
| OLFM1 | NR2F2 |
| PLXNA4 | PHACTR1 |
| PRKCQ | PXDC1 |
| SH3TC1 | RAB11FIP1 |
| SLC6A6 | RASL11B |
| | SEMA6A |
| | SLC1A4 |
| | TGFBI |
| | TIAM2 |

**Figure S4. BAZ1B knock down induces significant chromatin remodeling at distal regions.** (**A**) Regions differentially bound by BAZ1B in a WBS and 7dupASD *vs* CTL and atWBS comparison (FDR < 0.1, n=2). (**B**) Regions differentially bound by BAZ1B in a WBS *vs* 7dup, CTL and atWBS comparison (FDR < 0.1, n=2). (**C**) Regions differentially bound by BAZ1B in a 7dupASD *vs* CTL, WBS and atWBS comparison (FDR < 0.1, n=2). (**D**) Total number of genes that have their enhancers differentially marked following BAZ1B levels. (**E**) Overlap between genes whose expression follows BAZ1B levels (purple), genes whose enhancer-associated H3K27 acetylation follows BAZ1B levels (red) and genes bound by BAZ1B at their enhancers (blue). (**F**) Overlap between genes whose expression follows BAZ1B levels (purple), genes whose enhancer-associated H3K4 mono-methylation follows BAZ1B levels (red) and genes bound by BAZ1B at their enhancers (blue). (**G**) Overlap between genes whose expression follows BAZ1B levels (purple), genes whose enhancer-associated H3K27 trimethylation follows BAZ1B levels (red) and genes bound by BAZ1B at their enhancers (blue). (**H**) List of genes that, at the same time, follow BAZ1B levels, have their enhancers differentially marked concordantly (H3K27ac, H3K4me1 and H3K27me3) and are bound by BAZ1B at enhancers (overlap in Fig. 3G).
